# Large-Scale Genomic Analysis of CpG-Mediated Immunogenicity in Bacteriophages and a Novel Predictive Risk Index

**DOI:** 10.1101/2025.05.15.652987

**Authors:** Lukeman Kharrat, Camilo A. Garcia-Botero, Wade Ingersoll, Tiffany Luong, Alejandro Reyes, Dwayne R. Roach

**Affiliations:** Department of Biology, San Diego State University, San Diego, CA 92182, USA; Department of Biological Sciences, Universidad de los Andes, Bogotá, 111711, Colombia; Wellcome Sanger Institute Fellow, Hinxton, Cambridge, UK

**Keywords:** phage therapy, safety, pattern recognition, CpG motifs, immune response, host-phage interaction

## Abstract

Bacteriophage (phage) therapy is a promising alternative to antibiotics, yet phage-induced immune responses can affect treatment efficacy. However, current methods for assessing phage immunogenicity are limited, hindering the development of safer, more effective therapies. Here, we introduce the *Bacteriophage Risk Index* (BRI), a novel metric that quantifies phage immunogenic potential based on CpG dinucleotide frequency, motif spacing, and sequence context, key factors influencing Toll-like receptor 9 (TLR9) activation. Applying the BRI to 7,011 phage genomes, we classified them into five risk tiers, revealing substantial immunogenic variability, even among phages targeting the same bacterial host. BRI scores correlated with immune responses in human lung epithelial cells, validating its predictive power. Experimental testing further confirmed this, as exposure of lung epithelial cells to two phages from distinct risk tiers showed that the high-risk phage (Category 4) induced a strong pro-inflammatory response, upregulating CXCL1, CXCL8, IRF7, and TNFAIP3, while the low-risk phage (Category 2) triggered minimal immune activation with limited cytokine expression. These findings confirm that higher BRI scores predict stronger immune responses, providing a robust tool for evaluating phage immunogenicity. By enabling the selection of phages with lower immunogenic potential, the BRI enhances the safety and efficacy of phage therapy while offering a framework for regulatory agencies, clinical researchers, and biologic drug development, with applications extending beyond phage therapy to other immunogenic biologics.

## INTRODUCTION

The growing crisis of multidrug-resistant (MDR) bacterial infections poses a serious global health threat, with resistance to last-line antibiotics such as carbapenems and colistin becoming increasingly prevalent. In response, the World Health Organization (WHO) has classified MDR pathogens, including *Pseudomonas aeruginosa, Acinetobacter baumannii*, and *Enterobacteriaceae*, as priority threats, highlighting the urgent need for alternative therapeutic strategies (1). Among emerging approaches, bacteriophage (phage) therapy has gained renewed attention for its ability to selectively eliminate MDR pathogens while preserving the host microbiota, a significant advantage over conventional antibiotics (2-4). Phages not only circumvent common resistance mechanisms but also exhibit potent biofilm-degrading activity, addressing a major limitation of antibiotic therapy. Additionally, phage–antibiotic synergy has demonstrated the ability to enhance bacterial killing, restore antibiotic susceptibility, and reduce selective pressures that drive further resistance (2). This dual-action potential makes phage therapy a promising tool against refractory infections, with preclinical and clinical studies supporting its efficacy in cases where antibiotics alone have failed (5, 6). Indeed, several clinical trials are currently underway to systematically evaluate phage therapy for MDR infections, bolstering its translational potential and further validating its clinical utility (7, 8). However, despite its therapeutic promise, the successful clinical implementation of phage therapy is hindered by several key obstacles, with the most critical being the complex and variable interactions between phages and the host immune system (9-12).

Lytic double-stranded DNA (dsDNA) phages, particularly those within *Caudoviricetes*, are prime candidates for therapeutic applications due to their ability to rapidly infect and lyse bacterial hosts without integrating into the host genome (13). These viruses possess a distinctive tailed morphology that enables species-specific host recognition and facilitates efficient genome injection into bacterial cells. As the most diverse group of viruses, dsDNA phages exhibit extensive variation in genome size and content, which drive their adaptability and ecological success. Their genomes range from compact structures of approximately 10 kilobases (kb) to the expansive genomes exceeding 500 kb found in “jumbo” phages, reflecting remarkable evolutionary plasticity (14, 15). The combination of potent lytic activity, genetic stability, evolutionary adaptability, and target specificity highlights their ecological significance and expanding applications in both biotechnology and phage therapy (16, 17).

Phages, as foreign biological entities, inevitably engage the host immune system, provoking both innate and adaptive immune responses (9, 18, 19). While antibody-mediated neutralization is a well-documented challenge in phage therapy (20-23), innate immune recognition—particularly through pattern recognition receptors (PRRs)—remains an underappreciated yet critical barrier (9). Among these, unmethylated CpG motifs in phage DNA are recognized as microbe-associated molecular patterns (MAMPs), primarily activating Toll-like receptor 9 (TLR9) within endosomal compartments of phagocytes, B cells, and certain epithelial cells (e.g., in the gut, respiratory tract, or genitourinary tract). This mechanism is well-established in the recognition of bacterial and eukaryotic viral DNA (24-26).

Upon CpG binding, TLR9 undergoes a conformational change, recruiting the adaptor protein MyD88 and triggering downstream signaling cascades (27). These include the NF-κB pathway, which stimulates the production of pro-inflammatory cytokines (IL-6, TNF-α, and IFN-β), and the interferon regulatory factor (IRF) pathway, which induces type I interferons. While moderate immune activation can support bacterial clearance, excessive inflammation may accelerate phage elimination, thereby compromising therapeutic efficacy. Repeated exposure to phage DNA can also stimulate adaptive immunity, driving the production of neutralizing antibodies (IgG, IgA) that further inhibit phage–host interactions and impede bacterial targeting (28). Importantly, TLR9’s role in detecting exogenous DNA extends to other DNA-based therapeutics (e.g., plasmid DNA vaccines and gene therapy vectors), which may similarly face immune-mediated clearance and inflammatory responses (29). Balancing beneficial immune activation against the risk of excessive inflammation and neutralization is thus crucial for refining and advancing phage therapies and other DNA-based interventions alike.

Critically, the immunogenicity of phage DNA is not solely determined by the presence of CpG motifs, but also by their genomic context and arrangement. Factors such as CpG spacing, surrounding sequence features, and the presence of inhibitory sequence elements can significantly modulate TLR9 activation strength (30-32). While the precise role of CpG-mediated immunogenicity in phage therapy is still being elucidated, parallels from bacterial and viral studies suggest that optimizing phage selection may reduce unwanted immune responses. A comprehensive evaluation of these genomic and formulation factors will be essential for developing phage treatments with minimized immunogenicity, thereby enhancing both efficacy and safety in clinical applications.

In this study, we systematically analyzed 7,011 phage genomes associated with human-relevant bacterial hosts. We begin by characterizing the genomic dataset, including phage diversity and host distribution, and evaluate not only the presence of CpG motifs but also their genomic arrangement, encompassing CpG spacing, neighboring sequences, and inhibitory elements. In addition, we introduce a novel *Bacteriophage Risk Index* (BRI) to classify phages based on their likelihood of triggering TLR9, providing a new framework for assessing the immunological risk of using phages in therapeutic contexts. We further employed *in vitro* RNA-sequencing–based assays to elucidate how these features modulate TLR9-mediated immune responses, providing mechanistic insights that will guide the rational development of phages with minimized immunogenicity. As a immunogenicity assessment tool, the BRI holds promise for guiding phage therapy development, informing regulatory approval processes (FDA, EMA), and advancing personalized medicine.

## RESULTS

### Complete genome catalog of human-associated phages

We retrieved all complete genomes of dsDNA phages from NCBI’s Reference Sequence database on 12/04/2023. The dataset was filtered to include 7,011 genomes from dsDNA phages that infect one of 79 host species known to be associated with humans, including pathogens, commensals, foodborne bacteria, and probiotics (Figure 1a). The functional and genomic diversity of the phages in this dataset spans 18 of the 21 phage families, 209 genera, and several hundred species (Data File 1). Among the phage morphotypes, siphoviruses were the most prevalent, with 2,795 genomes, followed closely by myoviruses (2,667 genomes) and podoviruses (1,551 genomes). The dataset predominantly represented phages that infect host species in the phylum *Pseudomonadota* (formerly *Proteobacteria*), which accounted for approximately 70% of the strains, followed by phages infecting *Bacillota* (formerly *Firmicutes*) at around 27% (Figure 1a). Also represented were species in the phyla *Actinomycetota* (135), *Campylobacterota* (89), *Bdellovibrionota* (26), and *Bacteroidota* (4). As expected, the most represented host species are common targets of phage therapy, including *Enterobacteriaceae* spp. (3,188 genomes), *Pseudomonas aeruginosa* (684 genomes), and *Staphylococcus aureus* (419 genomes), and the widely used dairy production bacterium *Lactococcus lactis* (438 genomes).

**Figure 1.**
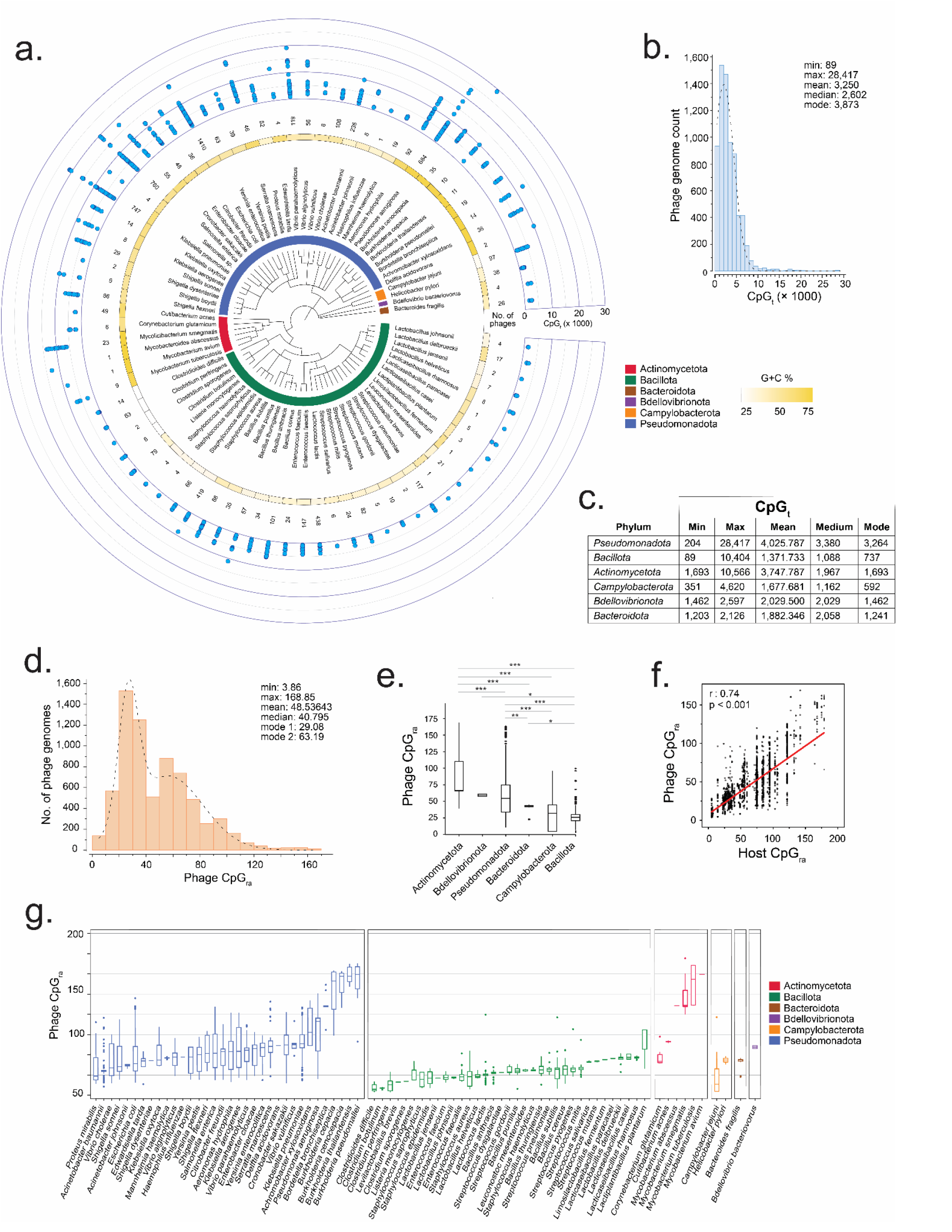
Diverse Distributions of CpG Motif Content and Relative Abundance in Phages Infecting Human-Relevant Bacterial Hosts. **(a)** Circular phylogenetic tree of 79 human associated host bacteria species and color-coded by host phylum (innermost ring). The second ring shows bacterial host G+C content, shading from light to dark yellow corresponding with low to high G+C content. The third ring represents the number of complete genomes of phages infecting human-relevant bacterial hosts from a dataset of 7,011 genomes. The outermost ring represents the total CpG dinucleotide content (CpGt) across each phage’s complete genome. **(b):** Histogram of CpGt values across all phages. The x-axis represents total CpG counts per genome (CpGt), while the y-axis indicates the number of phage genomes within each 100-motif interval. The distribution is right-skewed, with a peak at 3,873 CpG motifs. A subset of phage genomes contains exceptionally high CpG counts (>10,000), suggesting potential outliers with enhanced TLR9 activation potential. **(c)** Summary of the CpGt distribution by phylum, showing central tendency, spread, and distribution shape for phages grouped by their bacterial host phyla. **(d)** Histogram illustrating the distribution of CpG relative abundance per kilobase (CpGra) for all phages. The distribution is bimodal, with peaks at 29.08 and 63.19 CpGra. **(e)** Box-and-whisker plot showing the distribution of CpGra values across all phage genomes, stratified by bacterial host phylum. The plot displays the median, interquartile range, and outliers for each phylum. Statistical significance was assessed using one-way ANOVA with post hoc tests, with significance levels: ***p < 0.001, **p < 0.01, *p < 0.05. This plot highlights the variation in CpGra values among bacterial host phyla. **(f)** Scatter plot illustrating the correlation between phage CpGra and host CpGra. Each point represents a phage-host pair. A strong positive correlation was observed, with a Pearson’s correlation coefficient of r = 0.74 (p < 0.001), indicating a significant CpGra relationship between phage and host bacterial. **(g)** Box-and-whisker plot showing the distribution of CpGra values across all phage genomes, stratified by bacterial host species within each phylum. The plot displays the median, interquartile range (IQR), and outliers for each host species, ordered from lowest to highest CpGra within each host phylum. This plot illustrates the high variation in CpGra values among phage strains infecting different bacterial species. TLR9 activation, may reflect the dynamic interactions of their bacterial hosts with the human immune system, exemplified by species such as *Mycobacteria* spp. In contrast, *Bdellovibrionota* phages exhibit more uniform CpG content, as indicated by the lower mean CpGra (59.2 ± 2.2) and minimal variability. While *Bdellovibrionota* species primarily target other bacteria and indirectly influence the host immune system, the relatively consistent CpG content of their phages suggests a uniform immune activation potential. *Pseudomonadota* phages displayed a mean CpGra of 57.4 ± 26.2 with more moderate variability, suggesting an inconsistent immune activation potential across the group. Similar to *Actinomycetota* phages some *Pseudomonadota* phages may strongly trigger TLR9 activation while others, with lower CpGra, may evade immune detection. By comparison *Bacillota* and *Campylobacterota* phages, with the lowest CpGra values of 25.6 ± 9.0 and 28.5 ± 18.9 respectively, indicate a less prominent immune activation potential. The observed differences imply that both the phage’s environment, including human immune interactions, and the phage’s evolutionary adaptations to its bacterial host can play crucial roles in shaping CpG abundance. Indeed, a strong correlation (r = 0.74, p < 0.001) was found between the CpGra of phages and their corresponding host species (Figure 1f).

### CpG counts across phage genomes demonstrate vast TLR9 activation potentials

To assess the primary determinant of TLR9 activation, we first focused on the number of CpG motifs in a complete genome, as a higher number of these motifs increases the likelihood of TLR9 recognition and subsequent activation. We quantified the total number of CpG motifs (CpG_t_) in each phage genome within a dataset of 7,011 genomes (Figure 1a outer ring and Data File 2). Phage CpG_t_ values ranged from 89 to 28,417, with a mean of 3,250 and a median of 2,602, reflecting a positively skewed distribution (Figure 1b). This suggests that while most genomes have CpG counts concentrated towards the lower end of the range, a small subset contains CpG counts that are several times greater (e.g., >10,000). The most frequently observed CpG_t_ is 3,873, highlighting the commonality of a relatively moderate number of CpG in phage genomes.

Further comparison revealed substantial variability across phages within the same host phyla (Figure 1c). *Bacillota* phages, which infect bacteria such as *Staphylococcus* and *Clostridium*, exhibit the lowest mean CpG_t_ (1,371) but a broad content range (89–10,404), indicating high variability and heterogeneous CpG content compared to other groups. *Pseudomonadota* phages, infecting Gram-negative bacteria like the *Enterobacteriaceae* and *Pseudomonas*, have the highest mean CpG_t_ (4,025). Although their content range (204–28,417) is broader than *Bacillota*, their distribution is more concentrated around a mode of 3,264, indicating a skew towards moderate CpG content. *Actinomycetota* phages, which infect Gram-positives like *Mycobacterium* and *Corynebacterium*, show a high mean CpG_t_ (3,747), but their distribution has a narrower range (1,693–10,566) and is more concentrated at the lower end (mode = 1,693), suggesting less variation and a greater concentration of genomes with low CpG_t_. *Bacteroidota* and *Bdellovibrionota* phages exhibit a more concentrated CpG content, with ranges of 1,203–2,126 and 1,462–2,597, respectively, suggesting less variability compared to other phage groups. However, this finding likely also reflects their smaller sample sizes (Data File 2), suggesting that broader CpG_t_ ranges might emerge with additional phage discovery.

These findings suggest two key points: first, although CpG content in phage genomes varies significantly both across and within host phyla, it is more strongly correlated with their host species. Second, genomes with higher CpG motif frequencies, such as *Actinomycetota* phages, which tend to have high CpG counts, or certain outlier *Bacillota* phages with similarly elevated CpG counts, would be more likely to be detected by TLR9. For example, *Actinomycetota* and *Pseudomonadota* phages, which are rich in CpG motifs, would have up to four times the likelihood of TLR9 detection compared to other groups. In contrast, *Bacillota* phages, which have lower CpG frequencies, would be less likely to provoke an immune response.

### Widespread CpG motif content variability across phage genomes at the host species level

While CpG_t_ provides an initial measure of CpG motif abundance, direct comparisons across phage strains with varying genome sizes require normalization. To account for differences in genome length, we calculated CpG relative abundance per kilobase (CpG_ra_), allowing for a more standardized assessment of immunogenicity potential across host species. However, even after normalization considerable variability in CpG motif abundance remains, as reflected by the CpG_ra_ values ranging from 4 to 169 (Figure 1d vs Figure 1b). Therefore, genome size alone may not fully explain the observed differences in CpG content across the phages in the dataset. The distribution means of 49 and the slightly lower median of 41 suggest that most phage genomes are concentrated at the lower end of the spectrum (20–40 CpG_ra_) (Figure 1d and Data File 1). For context, human DNA has a CpG_ra_ of ∼24 (33). The distribution is largely bimodal, with two major peaks corresponding to CpG_ra_ of 29 and 63, respectively. This suggests that multiple factors such as phage type and host interaction may be contributing to the segregation of CpG abundance variability.

Host-specific trends contribute to the observed distributions in CpG abundance and immune activation potential. For example, *Actinomycetota* phages exhibited the highest mean CpG_ra_ (87.5 ± 33.6), indicating higher CpG densities with considerable variability between phage strains (Figure 1e). The diverse immune interaction potentials of these phages, which potentially increases the likelihood of

However, within each phylum, phages exhibit significant variability in CpG content across species, reflecting host-specific adaptations (Figure 1g). Even at the host species level phage CpG abundance remains variable, suggesting that factors beyond host-specific adaptations contribute to shaping CpG content and immune activation potential for individual phage strains. For instance, phages infecting *Burkholderia pseudomallei*, the causative agent of melioidosis, exhibit an exceptionally high CpG_ra_ mean (146.6 ± 14.6) with moderate-to-low variability (Figure 1g). In contrast, *Pseudomonas* phages show a moderately high CpG_ra_ mean (80.4 ± 91.2), with the highest degree of variability observed across the dataset. Phages infecting *Lactiplantibacillus plantarum*, a *Bacillota* species known for its broad digestive health and immune function benefits, also exhibit a moderately high CpG_ra_ mean (64.6 ± 27.3) with moderate variability. On the other hand, phages infecting species such as *Staphylococcus epidermidis* (21.2 ± 8.2), *Staphylococcus saprophyticus* (19.0 ± 10.6), *Enterococcus faecium* (22.0 ± 19.6), and *Campylobacter jejuni* (22.9 ± 19.6) display low mean abundances, yet exhibit significant variability in CpG content. Interestingly, some phages, such as those infecting *Cutibacterium* species (mean = 65.8 ± 0.6), show more uniform CpG_ra_ values. The overall observation is that phage CpG content varies significantly across species and phyla, reflecting both host-specific adaptations and other genomic factors influencing immune activation potential, with diverse phages infecting a single species showing high or low CpG content and variability, while others exhibit more conserved CpG patterns.

### CpG motif spacing and sequence context on TLR9 activation across phage genomes

The relationship between CpG content and TLR9 activation is not strictly linear. Rather, the interdistance between CpG motifs plays a crucial role in determining immune activation strength. We found that closer spacing between CpG motifs enhances TLR9 homodimer formation, triggering the conformational changes required for downstream signaling (34). Among 34 *Pseudomonadota* host species, 33 harbored phages where more than 50% of CpG pairs had a mean interdistance of <30 nucleotides (Figure 2a and Data File 3). The only exception was phages infecting *Acinetobacter baumannii*, an opportunistic pathogen with high multidrug resistance (MDR) and increasing global prevalence, which may reflect the host’s adaptation for immune evasion. Similarly, phages infecting *Actinomycetota, Bacteroidota*, and *Bdellovibrionota* generally exhibited closely spaced CpG motifs, with more than 50% of CpG pairs having a mean interdistance of <30 nucleotides. This suggests that phages from these species commonly enhance TLR9 homodimer formation, which likely strengthens immune activation. In contrast phages infecting *Bacillota* (29 of 35 species) and *Campylobacterota* (2 of 2 species) showed greater than 30 nucleotides between CpG pairs (Figure 2a and Data File 3). The larger interdistances between CpG motifs in these phages may impair TLR9 homodimer formation, potentially reducing immune activation efficiency. While many phages in the dataset have optimally spaced CpG motifs, phages infecting *Bacillota* and *Campylobacterota* species may have adapted to limit immune responses.

**Figure 2.**
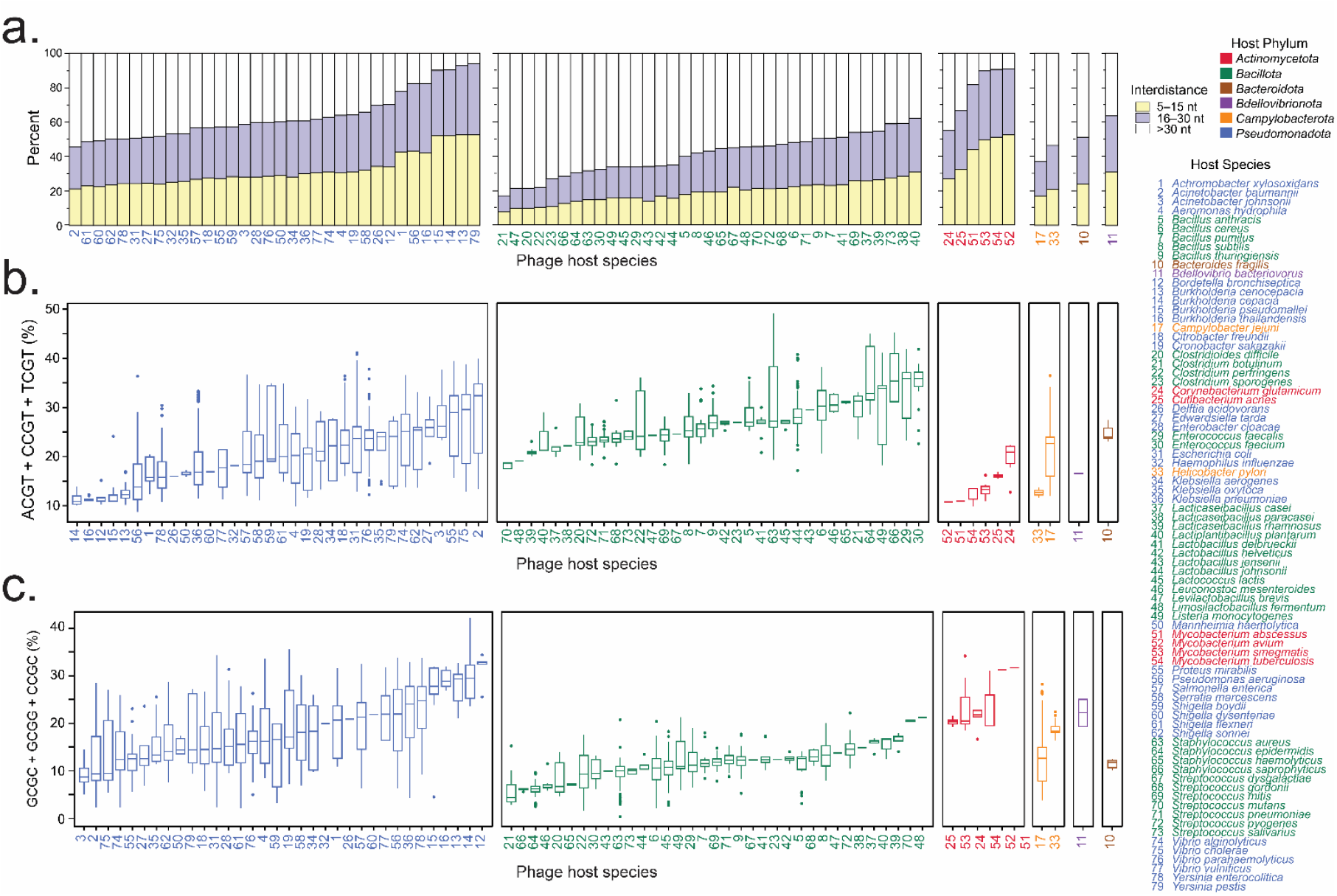
Phage genome CpG motif interdistances and the proportions within enhancing or inhibiting flanking sequences. **(a)** A stacked bar plot of CpG motif interdistance variability across phage genomes, grouped by bacterial host species and phylum, highlights potential differences in immune modulation. CpG pair interdistances are categorized into close (5-15 nucleotides; yellow), typically linked to amplified TLR9 activation, short (16-30 nucleotides; purple), generally associated with enhanced TLR9 activation, and long (>30 nucleotides; white), which may limit activation. The height of each bar represents the proportion of CpG pairs within each category. Phages are arranged along the x-axis based on their average CpG pair interdistance, ordered from shortest to longest within each bacterial host phylum. **(b and c)** Percentages of enhancing and inhibiting CpG motifs reflect potential functional differences in immune modulation across phage genomes. **(b)** The percentage of enhancing CpG motifs (ACGT, CCGT, TCGT) in phage genomes was calculated by summing their frequencies. **(c)** The percentage of inhibiting CpG motifs (GCGC, GCGG, CCGC) was calculated similarly. The plots display the median, interquartile range (IQR), and outliers for each host species, ordered from fewest to most CpG motifs in each tetramer collection within each host phylum. Phage host species are numbered alphabetically and colored by phyla: *Bacteroidota* (brown), *Bdellovibrionota* (purple), *Campylobacterota* (orange), *Pseudomonadota* (blue), *Actinomycetota* (red), and *Bacillota* (green).

Beyond CpG spacing, the specific tetramer sequences flanking CpG motifs also influence TLR9 binding, immune activation, and the stability of the TLR9-DNA complex (32). We analyzed the prevalence of tetramer sequences, such as ACGT, CCGT, and TCGT (enhancing), and GCGC, GCGG, and CCGC (inhibitory), which affect TLR9 recognition. Across phages grouped by host phyla, the distribution of CpG motifs in enhancing and inhibitory tetramers varied (Data File 4). *Pseudomonadota* phages, on average, had 20.1% of their CpG motifs in enhancing tetramers and 18.83% in inhibitory tetramers (Figure 2b, c), indicating a relatively balanced influence on TLR9 activation. Similarly, *Bacteroidota, Campylobacterota*, and *Bdellovibrionota* phages exhibited balanced distributions of CpG motifs in enhancing versus inhibitory tetramers, with frequencies of 24.71% vs. 22.19%, 17.14% vs. 18.64%, and 16.56% vs. 18.69%, respectively. In contrast, *Bacillota* phages had a higher proportion of CpG motifs in enhancing tetramers (26.93%) and a lower proportion in inhibitory tetramers (11.92%), suggesting a stronger potential for immune activation. This increased prevalence of enhancing tetramers could indicate an adaptation in *Bacillota* phages to compensate for their CpG-poor genomes (Figure 1), although the underlying mechanisms remain unclear. On the other hand, *Actinomycetota* phages possessed a higher proportion of CpG motifs in inhibitory tetramers (22.86%) compared to enhancing tetramers (13.82%) (Figure 2b, c), indicating a more inhibitory influence on immune activation despite their typically CpG-rich genomes (Figure 1).

Together, although many phages in the dataset exhibit optimally spaced CpG motifs, phages infecting *Bacillota* and *Campylobacterota* species may have evolved to limit immune detection. However, phages with CpG-poor genomes, like those infecting *Bacillota* species, may still induce strong immune responses due to a higher frequency of CpG motifs in enhancing tetramers. Our findings also emphasize that a higher overall CpG content, optimally spaced, would not necessarily correlate with stronger immune activation if a significant portion of those CpGs are part of inhibitory tetramers, as seen with *Actinomycetota* phages. Thus, the distribution of CpG motifs across enhancing and inhibitory tetramers could serve as a more accurate predictor of TLR9 activation and the overall immune response than CpG content alone.

### Assessing the full spectrum of phage immunogenicity risks

We observed significant variability in the immunogenic potential of phages, with each influencing factor, such as CpG content, spatial distribution and sequence patterns, demonstrating distinct levels of variation. There are few consistent patterns between these factors across all phage genomes, which prevents meaningful predictions about a phage strain’s immunogenic potential. To address this challenge, we developed the Bacteriophage Risk Index (BRI), a unified and normalized framework that combines these diverse elements into a single assessment score. The BRI categorizes phages into five immunological risk levels, ranging from minimal (Category 1) to critical (Category 5) (Figure 3). This structured risk framework provides precise safety assessments and offers actionable insights essential for tailoring phage therapeutics to ensure successful outcomes. For detailed equations, refer to the Methods.

**Figure 3.**
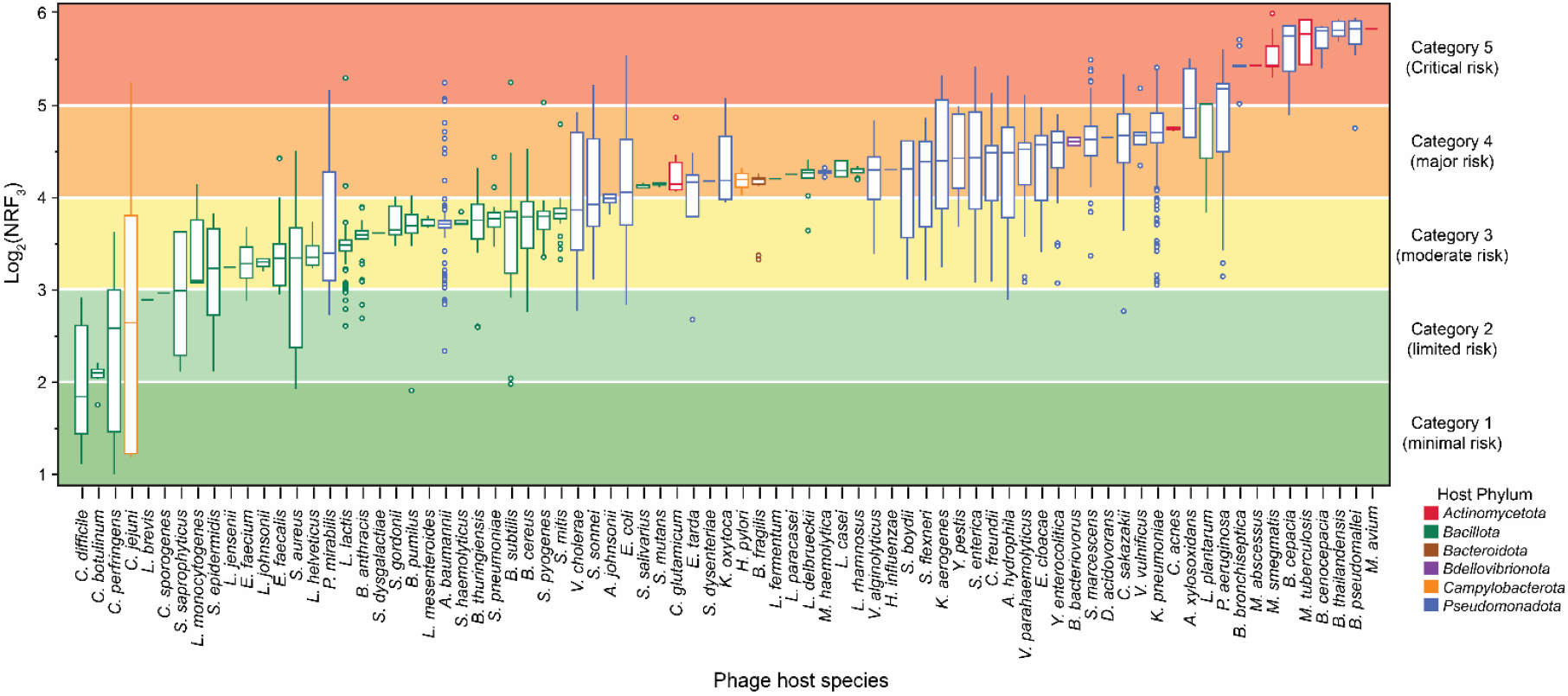
Bacteriophage Risk Index (BRI) harnesses phage immunogenicity heterogeneity for comparison within and across bacterial hosts. Box-and-whisker plot illustrates the distribution of normalized risk factor (NRF3) values for all phages infecting each host species, with bars colored by phyla: *Bacteroidota* (brown), *Bdellovibrionota* (purple), *Campylobacterota* (orange), *Pseudomonadota* (blue), Actinomycetota (red), and Bacillota (green). The plot displays the median, interquartile range (IQR), and outliers for each host species, arranged from lowest to highest NRF3 value. NRF3 values were classified into five BRI risk categories based on log2 transformations: Category 1 = minimal risk (1-2, dark green); Category 2 = limited risk (2-3, light green); Category 3 = moderate risk (3-4, yellow); Category 4 = major risk (4-5, orange); and Category 5 = critical risk (5-6, red). *(Note: higher resolution image at end of document)*

Phages in our dataset span all five risk categories, with 87% of genomes falling within the top three risk tiers (Figure 3). A small proportion (609 genomes) are classified into the critical risk Category 5, primarily consisting of phages infecting *Burkholderia, Mycobacterium*, and *Pseudomonas* species, which are significant health concerns due to their role in chronic, opportunistic, and frequent MDR infections. Category 4, representing major risk, includes 2,714 genomes that encompass phages targeting human pathogens associated with gastrointestinal infections, pneumonia, sexually transmitted infections, and foodborne illnesses. This category also includes phages targeting commensals such as *Bacteroides fragilis, Streptococcus mutans*, and *Lactobacillus plantarum*, along with common probiotic species like *Lactobacillus rhamnosus, Lactobacillus casei*, and *Lactobacillus delbrueckii*. Category 3 (2,780 genomes) contains phages that target a broad spectrum of pathogens and commensals, presenting moderate immunogenic potential. These species are linked to gastrointestinal, respiratory, skin, and soft tissue infections (e.g., *Vibrio cholerae, Streptococcus pneumoniae*, and *S. aureus*), with some hosts, such as *A. baumannii* and *Bacillus anthracis*, also causing severe systemic infections. Phages in this category also target members of the host microbiota, such as *Enterococcus faecalis, Streptococcus mitis*, and *Bacillus subtilis*, which can become opportunistic pathogens under certain conditions.

In lower-risk Categories 1 and 2, we classify the remaining 13% of phages (Figure 3). Although these phages pose a higher immunogenic risk than human “self” DNA, they primarily target species that, while part of the normal microbiota, can cause severe infections under certain conditions. For example, *C. jejuni, Clostridium perfringens*, and *Clostridium difficile* are generally harmless in healthy individuals but can become pathogenic under specific patient circumstances, highlighting the complex and context-dependent risk posed by phages infecting these species.

Additionally, we observe that individual phages infecting the same host species can span multiple risk categories depending on strain-specific characteristics (Data Files 2, 3 and 4). Of the 79 host species, 22 have phages spanning neighboring categories, while 14 host species span three risk categories (e.g., *P. aeruginosa, Shigella* spp., and *Streptococcus pyogenes*). Seven host species have phages spanning four categories (e.g., *Proteus mirabilis, Escherichia coli*, and *Aeromonas hydrophila*). *C. jejuni* phages span all five categories which is consistent with the variable nature of this host bacterium. This highlights the ability of the BRI to capture a full spectrum of immunogenic risk across a diverse range of phages with flexibility for granular risk classifications, allowing for more informed risk evaluations of phages that target both pathogenic and commensal bacteria.

### Contrasting immune profiles induced by low- and high-risk phages

We employed RNA sequencing to investigate gene expression changes in A549 human lung epithelial cells triggered by *Shigella* phage KPS64 (Category 2, BRI = 10) and *Pseudomonas* phage PAK_P1 (Category 4, BRI = 28). Our goal was to assess whether the BRI correlates with the scale and complexity of immune activation pathways, which represent limited and major immunological risks, respectively. Both phages induced significant changes in gene expression, with KPS64 and PAK_P1 triggering 477 and 531 differentially expressed genes (DEGs), respectively (Figure 4a, b). Despite the similar number of DEGs, the immune responses varied significantly in scope and intensity. PAK_P1 elicited a far more expansive and robust immune response (Figure 4c, Data File 6). KPS64 DNA caused modest upregulation of immune-related genes, such as IL32, TNFAIP3, and TNFRSF10C, involved in inflammation regulation and immune modulation. Additionally, chemokines like CXCL1 and CXCL2 were upregulated, promoting immune cell recruitment, while ICAM1 was elevated, indicating a role in immune cell trafficking. Antiviral responses were also suggested by the upregulation of IFIT2 and MX1, and IL27RA, which may fine-tune T cell activation. In contrast, PAK_P1 DNA triggered a far broader immune response, involving pathways related to chemokine signaling, immune cell recruitment, and antiviral defense (Figure 4c, Data File 6). Notable genes upregulated by PAK_P1 include CCL20, CCR7, CX3CL1, CXCL1, CXCL2, CXCL3, CXCL8, ICAM1, IRF7, PTX3, TNFAIP3, TNFRSF1B, TNFRSF6B, IFIT2, and IL27RA. This response indicates extensive immune recruitment and antiviral activation, with IRF7 stimulating interferon pathways and PTX3 driving the acute-phase response. The upregulation of TNFAIP3, TNFRSF1B, and TNFRSF6B suggest enhanced regulation of inflammation and immune activation, pointing to a more coordinated immune mobilization.

**Figure 4.**
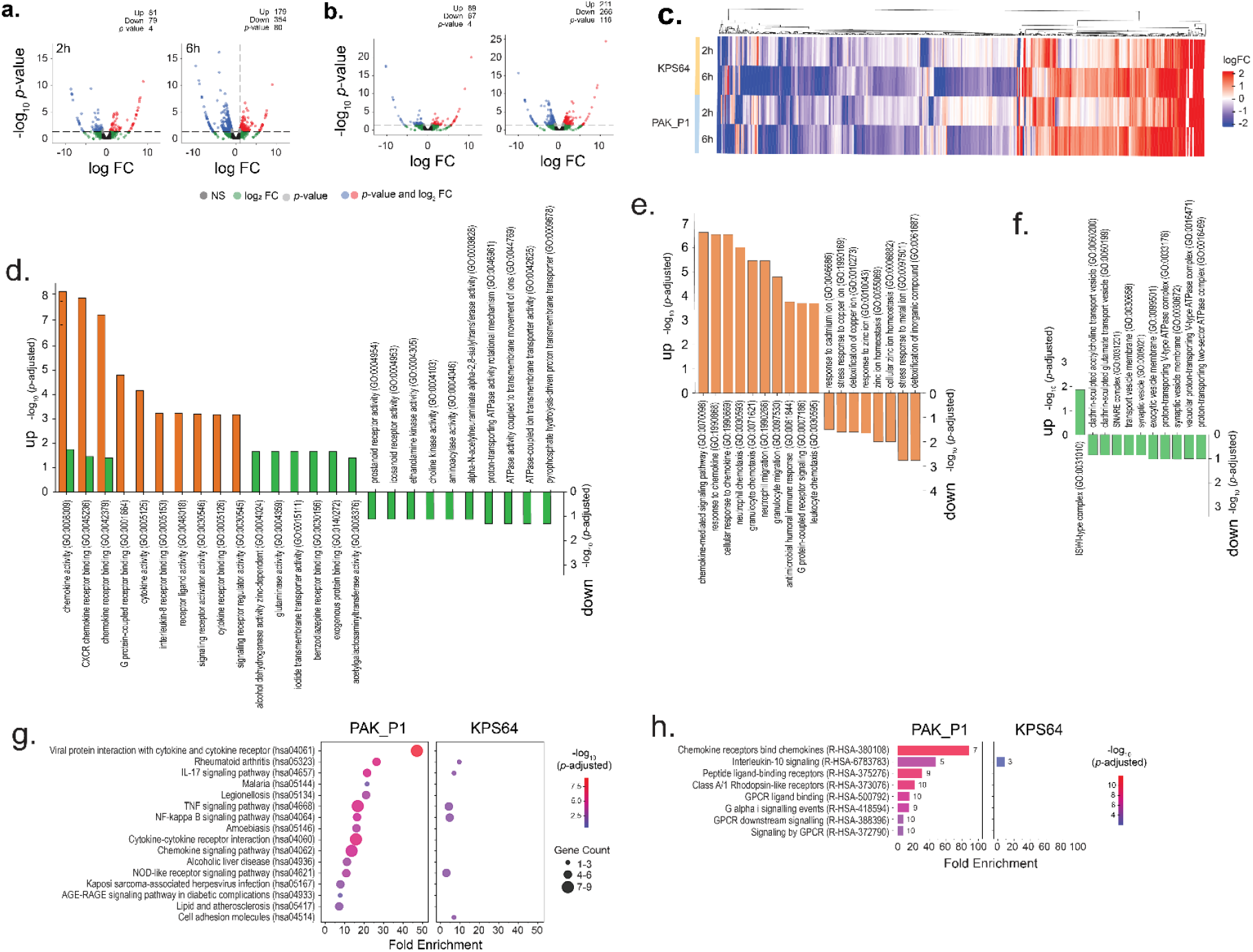
Transcriptomic analysis of differentially expressed genes (DEGs) in RNA sequencing data from A549 epithelial cells exposed to a Category 4 Pseudomonas phage PAK_P1 versus Category 2 Shigella phage KPS64. Volcano plot of pooled A549 epithelial cell DEGs (n=3) after 2 and 6 hours of exposure to **(a)** PAK_P1 and **(b)** KPS64. The x-axis represents the log2 fold change (FC), and the y-axis represents the -log10 of the p-value. Red points indicate significantly upregulated genes, and blue points represent significantly downregulated genes (p < 0.05, |FC| > 1). Green points are DEGs with log2FC > 1 but *p*-value >0.05. Non-significant genes are shown in black. **(c)** Heatmap of the expression patterns of the DEGs in A549 cells exposed to either PAK_P1 or KPS64. Rows represent samples, and columns represent individual genes. Color intensity indicates gene expression levels, with hierarchical clustering showing patterns of differential expression between the two phage exposures. **(d-f)** Gene Ontology (GO) enrichment analysis enrichment analysis for **(d)** biological processes, **(e)** cellular components, and **(f)** molecular functions after 6 hours. Significant GO terms (p < 0.05) are shown for each phage exposure. Bars for PAK_P1-exposed cells are shown in orange, while bars for KPS64-exposed cells are shown in green. The bar height corresponds to the enrichment score. **(g)** A dot plot depicting the top enriched Kyoto encyclopedia of genes and genomes (KEGG) pathways for A549 cells exposed to PAK_P1 and KPS64 for 6 hours. The size of the dots indicates the number of DEGs associated with each pathway, and the color represents the significance level (p-value). **(h)** Top Reactome pathways enriched in DEGs from A549 cells exposed to the two phages. The color gradient reflects the significance of each pathway (p < 0.05).

To further examine the immune responses triggered by KPS64 and PAK_P1 phage DNA, we conducted Gene Ontology (GO) analysis to identify relevant biological processes, alongside pathway enrichment analysis using KEGG and Reactome (Figure 4 d-g). The results revealed distinct immune profiles: KPS64 DNA induces a localized immune response, while PAK_P1 DNA triggers a broader and more coordinated immune activation. KPS64 DNA mainly activates immune cell recruitment and metabolic changes. Enriched GO terms such as chemokine activity (GO:0008009) and CXCR chemokine receptor binding (GO:0045236) suggest the immune response focuses on recruiting immune cells to the infection site (Figure 4d). Additionally, the activation of metabolic functions like alcohol dehydrogenase activity (GO:0004024) and glutaminase activity (GO:0004359) implies metabolic shifts supporting immune functions. However, the lack of enrichment in GO terms related to inflammation, antiviral defense, and pathogen clearance suggests a more restricted immune response. KEGG pathway analysis identified several immune-related pathways, including NF-kappa B signaling (Path:hsa04064), TNF signaling (Path:hsa04668), and NOD-like receptor signaling (Path:hsa04621), all involved in innate immunity, with some involvement of adaptive immunity via IL-17 signaling (Path:hsa04657) (Figure 4g). However, the response remains narrow. Downregulation of pathways linked to ion transport (GO:0042625, GO:0046961) and phospholipid synthesis (GO:0004103, GO:0004305) suggests cellular stress and disruptions in homeostasis.

In contrast, PAK_P1 DNA elicits a stronger, broader immune response. Pathway analysis reveals robust enrichment in pathways related to chemokine signaling, leukocyte migration, and antiviral defense. Key GO terms like neutrophil chemotaxis (GO:0030593, GO:0071621) and leukocyte migration (GO:0050900) suggest extensive immune cell recruitment (Figure 4d). Additionally, pathways linked to antimicrobial humoral immune responses (GO:0019730) and viral component response (GO:0071219) point to a potent antiviral defense. This extensive immune activation mirrors responses seen in infections caused by intracellular pathogens and viruses. PAK_P1 also activates a broader range of KEGG pathways, including Cytokine-Cytokine Receptor Interaction (Path:hsa04060) and Chemokine Signaling (Path:hsa04062), both of which enhance immune cell recruitment and intercellular communication (Figure 4g). Furthermore, NOD-like Receptor Signaling (Path:hsa04621) is more strongly activated, amplifying the immune response. Pathways related to intracellular pathogen recognition, such as Legionellosis (Path:hsa05134) and Kaposi Sarcoma-associated Herpesvirus Infection (Path:hsa05167), were also enriched, indicating a broader immune recognition compared to KPS64.

To further investigate the regulatory mechanisms, we analyzed Reactome pathways (Figure 4h). In response to KPS64 DNA, interleukin-10 signaling (R-HSA-6783783) was upregulated, indicating immune regulation to balance pro-inflammatory and anti-inflammatory responses. This suggests a localized immune response focused on inflammation regulation rather than broad activation. In contrast, PAK_P1 DNA also upregulated interleukin-10 signaling, but additionally triggered chemokine receptor binding (R-HSA-380108), enhancing immune cell recruitment, possibly driven by chemokines produced via TLR9 activation. The peptide ligand-binding receptors (R-HSA-375276) pathway was also enriched, suggesting a potential role for TLR9 in immune signaling. PAK_P1 DNA further activated GPCR signaling pathways, including G alpha signaling (R-HSA-418594) and GPCR downstream signaling (R-HSA-388396), highlighting the role of GPCRs in immune cell migration and cytokine release. The signaling by GPCR (R-HSA-372790) pathway emphasizes how chemokine receptors, potentially activated by TLR9-triggered chemokines, coordinate immune cell trafficking.

RNA sequencing of A549 human lung epithelial cells exposed to Shigella phage KPS64 (Category 2, BRI = 10) and Pseudomonas phage PAK_P1 (Category 4, BRI = 28) confirmed a strong correlation between BRI score and immune activation. KPS64 DNA induced a limited immune response, characterized by mild upregulation of cytokines such as IL-32 and TNFAIP3. In contrast, PAK_P1 triggered a broad immune activation profile, significantly upregulating chemokines (CXCL1, CXCL2, CXCL8) and inflammatory regulators (IRF7, TNFAIP3, ICAM1). An enrichment analysis revealed strong activation of the NF-kappa B and cytokine-cytokine receptor pathways in PAK_P1-exposed cells, consistent with a heightened immune response. These results validate the BRI as a predictive metric for CpG-mediated immunogenicity. It is important to note that this analysis captures immune responses at early time points (2 and 6 hours post-exposure), primarily reflecting acute-phase reactions. Future studies examining later time points are needed to assess potential shifts toward regulatory or adaptive immune responses, which may influence the long-term immunogenicity of phage DNA (9, 28).

## DISCUSSION

This study characterizes the immunogenic variability of phages and introduces the Bacteriophage Risk Index (BRI) as a predictive tool for assessing CpG-mediated immune activation. By analyzing 7,011 phage genomes, we identified key genomic features that may contribute to TLR9 activation, emphasizing the necessity for a systematic evaluation of immunogenicity in phage-based therapeutics. These findings align with prior research demonstrating that viral DNA activates innate immune responses through largely TLR9 recognition (24, 26). Importantly, the immunogenic potential of CpG motifs is determined not only by their abundance but also by motif spacing, sequence context, and the presence of inhibitory sequences, which may either amplify or suppress TLR9 activation (25, 32, 35). The BRI score variability observed among phages strains infecting the same bacterial host reinforces the complexity of human-phage interactions and underscores the necessity for precise, genome-informed phage immunogenicity assessments.

The biological relevance of the BRI was further supported by transcriptomic profiling of human lung epithelial cells, which revealed a strong correlation between BRI scores and immune response magnitude. Previous studies have demonstrated that phage-induced immune responses are highly variable, influenced by genomic composition and host factors (20, 36, 37). In our study, phages classified as high-risk, such as *Pseudomonas* phage PAK_P1 (BRI = 28), induced significant upregulation of pro-inflammatory cytokines (CXCL1, CXCL2, CXCL8) and interferon regulatory factors (IRF7). This response is consistent with findings demonstrating that CpG-rich viral DNA enhances inflammatory signaling via TLR9-mediated activation of NF-kappa B and cytokine-cytokine receptor interactions (25, 38). Conversely, low-risk phages, including *Shigella* phage KPS64 (BRI = 10), elicited minimal cytokine expression, reinforcing prior observations that certain phages are tolerated by the immune system (39). However, additional experimental validation using direct immune assays is required to fully elucidate these mechanisms and determine whether phage structural proteins also contribute to immune recognition (40, 41).

The variability in phage-induced immune responses underscores the need for individualized assessments in therapeutic applications. While phages are generally recognized as safe (GRAS), recent clinical findings indicate that highly immunogenic strains may exacerbate inflammatory responses, particularly in patients with cystic fibrosis, inflammatory bowel disease, or autoimmune disorders (11, 21). These findings highlight the necessity of integrating immune profiling into phage selection, ensuring that therapeutic candidates align with individual patient immune status. Future studies should investigate how host-specific factors, including genetic predisposition, prior microbial exposures, and immune status, influence responses to CpG-rich phages. To address these challenges, the BRI offers a structured framework for assessing phage immunogenicity, enabling the selection of therapeutic candidates with lower immune activation potential. By incorporating BRI scores into phage selection workflows, clinicians could design personalized phage cocktails that optimize antibacterial efficacy while minimizing immune-related adverse effects. Additionally, integrating BRI assessments with patient-specific immune profiling could advance a precision-medicine approach to phage therapy, ensuring treatments are tailored to individual immune responses.

Beyond phage therapy, the principles underlying the BRI, particularly its assessment of CpG motif patterns, may have broader applications in biologic therapeutics, including gene therapy vectors and vaccines. Wright (32) demonstrated that CpG motifs in recombinant AAV gene therapy vectors contribute to TLR9-mediated immune activation, underscoring the potential relevance of BRI-like risk assessment models in biologic safety evaluations. Furthermore, the BRI framework could be adapted to assess oncolytic virotherapies and viral-vector-based immunotherapies, where immune activation is a key determinant of both efficacy and safety.

A critical next step in optimizing phage-based and biologic therapies is the standardization of immunogenicity assessments within regulatory frameworks. Current biologic safety assessments primarily focus on known genetic risk factors, such as toxin genes, antibiotic resistance elements, and lysogeny markers (42, 43). However, these methods fail to capture the full spectrum of host immune interactions with phage DNA. Integrating predictive risk assessment models such as the BRI into regulatory frameworks, including those established by the FDA and EMA, could enhance the standardization of phage therapy approvals, ensuring more comprehensive immunogenicity evaluations and facilitating clinical translation.

Despite the advances made in this study, several limitations must be acknowledged. The transcriptomic analysis captured only early-phase immune responses (up to 6 hours post-exposure), primarily reflecting acute inflammatory reactions. Longitudinal studies assessing immune dynamics over extended time periods will be necessary to determine whether high-BRI phages induce sustained inflammation or transition toward regulatory immune responses. Additionally, in vivo validation using mammalian models is required to assess systemic immune modulation by phage DNA and its implications for therapeutic outcomes. Moreover, this study focused on phage DNA immunogenicity, but future research should investigate the role of phage structural proteins in modulating immune recognition and activation. Previous studies have shown that capsid proteins and tail fibers can engage pattern recognition receptors (PRRs), contributing to immune activation (40, 44). Moreover, contaminants in phage preparations, including lipopolysaccharides (LPS) and bacterial debris, can confound immunogenicity assessments, emphasizing the need for rigorous purification protocols (45).

In conclusion, this study presents the BRI as a novel and systematic metric for predicting phage immunogenicity, addressing a critical gap in phage therapy safety assessments. By quantifying CpG-mediated immune activation, the BRI provides a framework for selecting therapeutic phages with optimized immunogenic profiles, enhancing both safety and efficacy in phage-based treatments. As phage therapy advances toward clinical integration, predictive risk assessment models like the BRI will be essential for developing precision-based therapeutics, mitigating immune-related risks, and streamlining regulatory approvals. Expanding the BRI framework to include epigenetic modifications and phage structural proteins will further refine its predictive accuracy and extend its application to viral vector-based gene therapies, oncolytic virotherapies, and vaccine development. Standardized immunogenicity assessments for phage-based and biologic therapeutics will be crucial for ensuring safety, optimizing efficacy, and facilitating regulatory acceptance, ultimately advancing the next generation of precision-engineered biologics.

## METHODS

### Bacterial host genome selection

A systematic search was performed across the NCBI BioProject (https://www.ncbi.nlm.nih.gov/bioproject) and PubMed (https://pubmed.ncbi.nlm.nih.gov/) databases to identify well-documented bacterial species with known human associations, categorized as pathogens, pathobionts, foodborne agents, zoonotic bacteria, commensals, or probiotics. Bacterial species without an associated phage genome in NCBI RefSeq database (https://www.ncbi.nlm.nih.gov/refseq/) were excluded from further analysis. After manual curation and review, a final list of 79 bacterial species was compiled, each associated with at least one phage genome.

### Bacterial strains, phages

*Pseudomonas aeruginosa* strains PAK (NZ_CP020659.1) and *Shigella flexneri* PE577 (CP042980.1) were used as laboratory strains to produce phage lysates. Host strains were grown overnight in Luria Broth (LB) Lennox medium at 37°C with shaking.

*Pseudomonas* phage PAK_P1 (GCF_000891855.1) and *Shigella* phage KPS64 (MK562502.1), both myoviruses, have genomes of 93.2 kb and 90.2 kb, respectively. Phage strains were confirmed free of toxins, antibiotic resistance genes, and lysogeny-related elements. Enrichment was performed with an MOI of 0.1, yielding 50mL lysates, which were sterilized through double centrifugation and filtration at a 0.22 μm cutoff. Phage DNA was extracted using the phenol-chloroform method to maintain quality and prevent shearing. Bacterial lysates were treated with DNase, Proteinase K, and 10% SDS, followed by incubation at 55°C for 30 min. The lysate was then mixed with equal volumes of phenol:chloroform:isoamyl alcohol in Phase Lock Gel tubes and centrifuged at 12,000 x g for 10 min to separate the phases. The aqueous phase containing DNA was transferred to a fresh tube and re-extracted with fresh phenol:chloroform:isoamyl alcohol. A final extraction with 100% chloroform removed phenol contamination. DNA was precipitated with 3M sodium acetate (pH 5.2) and ice-cold ethanol, incubated at -20°C for 30 min, and centrifuged at 12,000 x g for 20 min at 4°C. The DNA pellet was washed with 70% ethanol, air-dried, and resuspended in molecular-grade water. The final DNA preparation was suitable for downstream applications such as PCR and sequencing.

### Bacterial phylogeny

To construct a phylogenetic tree for the host species, taxonomy IDs were retrieved from the NCBI Taxonomy database (https://www.ncbi.nlm.nih.gov/taxonomy) and used in *PhyloT* v2 (https://phylot.biobyte.de/) to construct a Newick file. These data were subsequently integrated into the phylogenetic tree visualization in *R* using the *ggtreeExtra* package (v. 1.12.0) with default setting (v. 1.12.0) (46).

### Phage genome selection

All complete phage genome sequences were retrieved from NCBI’s RefSeq database (accessed on 12/04/2023) and first filtered for dsDNA phages belonging to the *Caudoviricetes* class, and then the aforementioned 79 human-associated bacterial species. Genomes described as “fragment” or “partial” were excluded. A custom Python script was used to extract phage’s host taxonomic information and phage taxonomic data. Phages lacking specific taxonomic classifications were grouped as “Unclassified.” The final dataset is provided in Data File 1.

### Nucleotide k-mer counts and relative abundance

To determine k-mer frequencies and relative abundances in complete phage genomes, we employed custom Python and Perl scripts to analyze FASTA-formatted sequences. Sequences were first parsed to exclude non-standard nucleotides (i.e., bases other than A, T, C, and G), and missing data were removed to ensure accurate analysis. Ambiguous sequences were accounted for using IUPAC nucleotide codes.

The occurrences of mono-, di-, tri-, and tetranucleotides, as well as genome lengths and G+C content, with counts performed on both forward and reverse strands were computed. K-mer counts were normalized by averaging the frequency per kilobase (kb) of DNA, yielding relative abundances; in particular, the normalized CpG frequency (CpG_ra_) representing the number of CpG dinucleotides per kilobase. Total CpG (CpG_t_) was calculated by multiplying the percentage frequency of CpG by the total genome size in base pairs. To efficiently manage large datasets, oversized input files were automatically partitioned into smaller fragments based on predefined memory constraints. This data is provided in Data File 2.

### CpG interdistance

The empirical inter-CpG distances for non-overlapping CGN[X]CG sites were calculated, where N[X] represents any nucleotide (A, C, G, or T) in the sequence between two CpG sites. A custom Python script processed the FASTA-formatted sequences, scanning both strands to identify CpG sites. For each genome, the script located consecutive, non-overlapping CpG sites and measured the number of nucleotides separating them. The resulting distances were grouped into three bins: 5–15 nt, 16–30 nt, and >30 nt. These binned results are available in Data File 3.

### TLR9 activation potential

To quantify the TLR9 activation potential of phage DNA, we adapted an approach developed by Wright to assess eukaryotic viruses (32) to incorporate more key genomic features well-known known to influence TLR9 activation. This modified method evaluates the immunogenicity of individual phage CpG-DNA sequences through a series of progressively refined risk factor (RF) equations. The first equation, RF_1_, calculates the relative abundance of total CpG motifs (CpGt) in the genome, normalized by the genome length in nucleotides (nt):

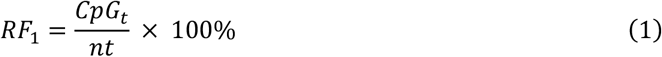

The second equation, RF_2_, builds upon RF1 by incorporating the presence of stimulatory ACGT, TCGT, and CCGT and inhibitory GCGC, GCGG, and CCGC motif sequences. In RF2, each motif type is assigned an activation weight: +1 for stimulatory motifs (S_4_) and -2 for inhibitory motifs (I_4_):

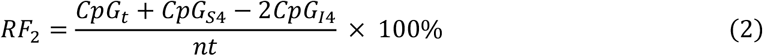

The third equation, RF_3_, further refines the model by incorporating CpG inter-distance, which represents the spacing between consecutive CpG motifs. Empirical studies suggest that shorter inter-distances enhance TLR9 activation (31). Activation weights are assigned as follows: a weight of +2 for distances between 5–15 nt (CpG5−15) and a weight of +1 for distances between 16–30 nucleotides (CpG16−30):

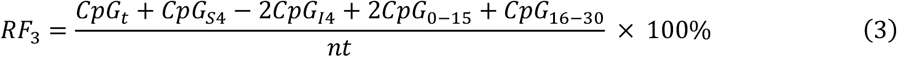

To normalize the results, NRF_3_ values were then scaled to a human genome reference value of 0.191, which represents the baseline or minimal TLR9 activation potential:

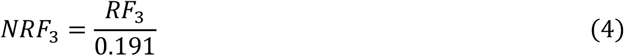

To improve interpretability and account for the skewed distribution of NRF3 values, we applied a log2 transformation. This approach ensures that immunogenicity risk is categorized based on relative rather than absolute differences, preventing high-CpG outliers from disproportionately affecting risk classification. Category 1 (log2 NRF_3_ values between 1 and 2) estimates minimal risk; Category 2 (log2 NRF_3_ values between 2 and 3) estimates limited risk; Category 3 (log2 NRF_3_ values between 3 and 4) estimates moderate risk; Category 4 (log2 NRF_3_ values between 4 and 5) estimates major risk; and Category 5 (log2 NRF_3_ values between 5 and 6) estimates critical risk. Scores are listed in Data File 5.

### Cell culture preparation

The lung epithelial cell line A549 was obtained from ATCC (Manassas, VA, USA) and cultured in Dulbecco’s Modified Eagle Medium (DMEM) supplemented with 10% fetal bovine serum (Gibco) at 37°C in 5% CO_2_. A549 cells were seeded at 8×10^5^ cells/well and grown for 24h to reach a density of 1×10^6^ cells/well in a 6-well tissue culture plate. After 24 hours of acclimation in serum-free DMEM, fresh serum-free DMEM was given to the cells, and they were treated for 2h or 6h with 10 μg/mL of purified DNA. Control A549 cells were cultured under the same conditions but incubated with sterile Dulbecco’s phosphate-buffered saline (dPBS) for 2h.

### RNA collection and sequencing

Total RNA was isolated from cells using the NucleoSpin RNA Plus isolation kit (Macherey-Nagel). The RNA was then reverse-transcribed into cDNA using the High-Capacity cDNA Reverse Transcription Kit with RNase inhibitor (Applied Biosystems), following the manufacturer’s instructions. The cDNA samples were stored at -80°C until further use. RNA sequencing was conducted at MiGS (Pittsburgh) using the NovaSeq platform (Illumina), adhering to the standard manufacturer’s protocols.

Approximately 50 million 150 bp paired-end reads were generated for each sample. Quality control and adapter trimming were performed using Illumina’s *bcl2fastq2* Conversion Software (v. 2.20). Read quantification was carried out with RSEM (47), and read counts were normalized using *edgeR’s* (48) Trimmed Mean of M values (TMM) algorithm. The resulting values were then converted to counts per million (cpm).

Differential expression analysis was conducted using *edgeR’s* exact test (exactTest) functionality between treatment groups. Given the absence of replicates in each group, dispersion was estimated at 0.04, as recommended by the *edgeR* user manual. Genes with an absolute log fold change |logFC| > 1 and a p-value < 0.05 were considered differentially expressed.

### Differential gene expression analysis

Differentially expressed genes (DEGs) were identified using *edgeR*’s exact test (*exactTest*) between treatment groups. As biological replicates were sequenced together in a single group, dispersion was set to 0.04, as recommended by the *edgeR* user manual. Genes with an absolute log fold change (|logFC|) > 1 and a p-value < 0.05 were classified as differentially expressed. Additionally, significance was further evaluated using a false discovery rate (FDR) threshold of < 0.05. DEGs identified using *edgeR* were visualized with volcano plots generated via the *EnhancedVolcano* function from the *Bioconductor* package (v. 3.18) in *R*. Additionally, heatmaps were produced to illustrate the expression patterns of these DEGs. For the heatmaps, the data were first converted into matrix format and then plotted using the *ComplexHeatmap* function from *Bioconductor* (v. 3.18). Together, these visualizations display the DEGs for both PAK_P1 and KPS64 at 2-and 6-hours following exposure of A549 epithelial cells to purified phage DNA.

### Functional enrichment analyses

Gene ontology (GO), KEGG, and Reactome pathway enrichment analyses were conducted using *R* packages *clusterProfiler* and *ReactomePA* from *Bioconductor* (v. 3.18). Gene symbols were mapped to Entrez IDs using *AnnotationDbi* and *org*.*Hs*.*eg*.*db* from *Bioconductor* (v. 3.18). Enrichment analyses were performed with *enrichGO* (for GO terms), *enrichKEGG* (for KEGG pathways), and *enrichPathway*, using all quantified genes as the background. Enriched terms with an p-adj < 0.05 were considered significant, and the top pathways for each category were curated to reduce redundancy.

### Statistical Analyses

All numerical data are presented as raw counts, mean ± standard deviation (SD), or percentage (%), as appropriate. Statistical analyses were conducted using R Studio software with default settings, unless otherwise specified. To assess correlations, the Pearson correlation coefficient (r) was used. Statistical significance was defined as a p-value < 0.05.

## Code availability

Data were analyzed and processed using R Studio, Python, Jupyter Notebook, Perl, and Bash scripting within a standardized pipeline, unless otherwise specified. Custom scripts used in the analysis are available upon request.

## Acknowledgments

This work was supported by the NIH National Institute of Allergy and Infection (Award Number R01AI177997) and The Conrad Prebys Foundation.

## Author Contributions

Conceptualization, DR, LK, AR; methodology, LK, DR, CAG, AR; code, LK, CAG, AR; investigation, LK, DR, AR, WI, TL; formal analysis, LK, DR; images, LK, DR; writing and editing, LK, TL, AR, DR.

## List of Supplementary Data

Data Files 1 and 2 supporting Figure 1.

Data Files 3 and 4 supporting Figure 2.

Data Files 5 supporting Figure 3.

Data Files 6 and 7 supporting Figure 4.

RNA seq: GEO accession GSE293045

## REFERENCES

1. S. Bertagnolio et al., WHO global research priorities for antimicrobial resistance in human health. The Lancet Microbe 5 (2024).

2. T. Luong, A. C. Salabarria, D. R. Roach, Phage therapy in the resistance era: Where do we stand and where are we going? Clin Ther 42, 1659–1680 (2020).

3. G. A. Suh et al., Considerations for the use of phage therapy in clinical practice. Antimicrob Agents Chemother 66, e0207121 (2022).

4. S. A. Strathdee, G. F. Hatfull, V. K. Mutalik, R. T. Schooley, Phage therapy: From biological mechanisms to future directions. Cell 186, 17–31 (2023).

5. G. A. Suh et al., Considerations for the use of phage therapy in clinical practice. Antimicrobial Agents and Chemotherapy 66, e02071–02021 (2022).

6. S. Uyttebroek et al., Safety and efficacy of phage therapy in difficult-to-treat infections: a systematic review. The Lancet Infectious Diseases 22, e208–e220 (2022).

7. P. D. Tamma et al., Safety and microbiological activity of phage therapy in persons with cystic fibrosis colonized with Pseudomonas aeruginosa: study protocol for a phase 1b/2, multicenter, randomized, double-blind, placebo-controlled trial. Trials 23, 1057 (2022).

8. K. Diallo, A. Dublanchet, A century of clinical use of phages: A literature review. Antibiotics (Basel) 12 (2023).

9. K. Champagne-Jorgensen, T. Luong, T. Darby, D. R. Roach, Immunogenicity of bacteriophages. Trends Microbiol 31, 1058–1071 (2023).

10. M. Lusiak-Szelachowska, R. Miedzybrodzki, W. Fortuna, J. Borysowski, A. Gorski, Anti-phage serum antibody responses and the outcome of phage therapy. Folia Microbiol (Praha) 66, 127–131 (2021).

11. S. Aslam et al., Pseudomonas aeruginosa ventricular assist device infections: findings from ineffective phage therapies in five cases. Antimicrob Agents Chemother 68, e0172823 (2024).

12. S. C. Nang et al., Pharmacokinetics/pharmacodynamics of phage therapy: a major hurdle to clinical translation. Clin Microbiol Infect 29, 702–709 (2023).

13. N. M. Smith et al., A mechanism-based pathway toward administering highly active N-phage cocktails. Front Microbiol 14, 1292618 (2023).

14. H. Brüssow, R. W. Hendrix, Phage genomics: small is beautiful. Cell 108, 13–16 (2002).

15. K. R. Harding, N. Kyte, P. C. Fineran, Jumbo phages. Curr Biol 33, R750–r751 (2023).

16. C. R. de Vries et al., Phages in vaccine design and immunity; mechanisms and mysteries. Current opinion in biotechnology 68, 160–165 (2021).

17. L. Gay, K. Suwan, A. Hajitou, Construction and utilization of a new generation of bacteriophage-based particles, or TPA, for guided systemic delivery of nucleic acids to tumors. Nature Protocols 10.1038/s41596-024-01040-9 (2024).

18. K. Gembara, K. DÁbrowska, Phage-specific antibodies. Current opinion in biotechnology 68, 186–192 (2021).

19. D. R. Roach et al., Human neutrophil response to Pseudomonas bacteriophage PAK_P1, a therapeutic candidate. Viruses 15 (2023).

20. R. M. Dedrick et al., Potent antibody-mediated neutralization limits bacteriophage treatment of a pulmonary Mycobacterium abscessus infection. Nat Med 27, 1357–1361 (2021).

21. M. Bernabeu-Gimeno et al., Neutralizing antibodies after nebulized phage therapy in cystic fibrosis patients. Med 5, 1096–1111 e1096 (2024).

22. J. D. Berkson et al., Phage-specific immunity impairs efficacy of bacteriophage targeting Vancomycin Resistant Enterococcus in a murine model. Nat Commun 15, 2993 (2024).

23. O. Krut, I. Bekeredjian-Ding, Contribution of the Immune Response to Phage Therapy. J Immunol 200, 3037–3044 (2018).

24. J. Pohar et al., Selectivity of human TLR9 for double CpG motifs and implications for the recognition of genomic DNA. J Immunol 198, 2093–2104 (2017).

25. A. Dalpke, J. Frank, M. Peter, K. Heeg, Activation of toll-like receptor 9 by DNA from different bacterial species. Infect Immun 74, 940–946 (2006).

26. S. Bauer et al., Human TLR9 confers responsiveness to bacterial DNA via species-specific CpG motif recognition. Proc Natl Acad Sci U S A 98, 9237–9242 (2001).

27. B. Briard, D. E. Place, T. D. Kanneganti, DNA Sensing in the Innate Immune Response. Physiology (Bethesda) 35, 112–124 (2020).

28. M. Podlacha et al., Bacteriophage DNA induces an interrupted immune response during phage therapy in a chicken model. Nat Commun 15, 2274 (2024).

29. R. Sartorius, M. Trovato, R. Manco, L. D’Apice, P. De Berardinis, Exploiting viral sensing mediated by Toll-like receptors to design innovative vaccines. NPJ Vaccines 6, 127 (2021).

30. L. Han, B. Su, W. H. Li, Z. Zhao, CpG island density and its correlations with genomic features in mammalian genomes. Genome Biol 9, R79 (2008).

31. J. Pohar, A. Kuznik Krajnik, R. Jerala, M. Bencina, Minimal sequence requirements for oligodeoxyribonucleotides activating human TLR9. J Immunol 194, 3901–3908 (2015).

32. J. F. Wright, Quantification of CpG motifs in rAAV genomes: avoiding the toll. Mol Ther 28, 1756–1758 (2020).

33. D. Takai, P. A. Jones, Comprehensive analysis of CpG islands in human chromosomes 21 and 22. Proc Natl Acad Sci U S A 99, 3740–3745 (2002).

34. U. Ohto et al., Toll-like receptor 9 contains two DNA binding sites that function cooperatively to promote receptor dimerization and activation. Immunity 48, 649–658 e644 (2018).

35. D. Bayik, I. Gursel, D. M. Klinman, Structure, mechanism and therapeutic utility of immunosuppressive oligonucleotides. Pharmacol Res 105, 216–225 (2016).

36. M. Popescu, J. D. Van Belleghem, A. Khosravi, P. L. Bollyky, Bacteriophages and the immune system. Annu Rev Virol 8, 415–435 (2021).

37. M. Lusiak-Szelachowska et al., Phage neutralization by sera of patients receiving phage therapy. Viral Immunol 27, 295–304 (2014).

38. P. Knuefermann et al., CpG oligonucleotide activates Toll-like receptor 9 and causes lung inflammation in vivo. Respir Res 8, 72 (2007).

39. J.-P. Pirnay et al., Personalized bacteriophage therapy outcomes for 100 consecutive cases: a multicentre, multinational, retrospective observational study. Nature Microbiology 9, 1434–1453 (2024).

40. K. Dabrowska et al., Hoc protein regulates the biological effects of T4 phage in mammals. Archives of microbiology 187, 489–498 (2007).

41. K. Dabrowska et al., Immunogenicity studies of proteins forming the T4 phage head surface. J Virol 88, 12551–12557 (2014).

42. J. Y. Nale, M. R. J. Clokie, Preclinical data and safety assessment of phage therapy in humans. Current opinion in biotechnology 68, 310–317 (2021).

43. M. Koncz et al., Genomic surveillance as a scalable framework for precision phage therapy against antibiotic-resistant pathogens. Cell 187, 5901-5918.e5928 (2024).

44. K. Dabrowska, K. Switala-Jelen, A. Opolski, B. Weber-Dabrowska, A. Gorski, Bacteriophage penetration in vertebrates. J Appl Microbiol 98, 7–13 (2005).

45. T. Luong, A.-C. Salabarria, R. A. Edwards, D. R. Roach, Standardized bacteriophage purification for personalized phage therapy. Nature Protocols 15, 2867–2890 (2020).

46. S. Xu et al., ggtreeExtra: compact visualization of richly annotated phylogenetic data. Molecular Biology and Evolution 38, 4039–4042 (2021).

47. B. Li, C. N. Dewey, RSEM: accurate transcript quantification from RNA-Seq data with or without a reference genome. BMC Bioinformatics 12, 323 (2011).

48. M. D. Robinson, D. J. McCarthy, G. K. Smyth, edgeR: a Bioconductor package for differential expression analysis of digital gene expression data. Bioinformatics 26, 139–140 (2010).

